# scGFT: single-cell RNA-seq data augmentation using generative Fourier transformer

**DOI:** 10.1101/2024.07.09.602768

**Authors:** Nima Nouri

## Abstract

Integrating single-cell RNA sequencing (scRNA-seq) with artificial intelligence (AI) ushers in a new frontier for advanced therapeutic discoveries. However, for this synergy to achieve its full potential, extensive datasets are required to effectively train the AI component. This demand is particularly challenging when delving into rare diseases and uncommon cell types. Generative models designed to address data scarcity often face similar limitations due to their reliance on pre-training, inadvertently perpetuating a cycle of data inadequacy. To overcome this obstacle, we introduce scGFT (single-cell Generative Fourier Transformer), a train-free, cell-centric generative model adept at synthesizing single cells that exhibit natural gene expression profiles present within authentic datasets. Using both simulated and experimental data, we demonstrate the mathematical rigor of scGFT and validate its ability to synthesize cells that preserve the intrinsic characteristics delineated in scRNA-seq data. By streamlining single-cell data augmentation, scGFT offers a scalable solution to overcome data scarcity and holds the potential to advance AI-driven precision medicine.

## Main

The application of single-cell RNA sequencing (scRNA-seq) has become increasingly prominent in research settings, notably for its pivotal role in elucidating disease mechanisms and treatment responses at the cellular level^1^. Simultaneously, advancements in deep learning and artificial intelligence (AI) are revolutionizing medicinal applications, particularly in drug discovery^2-4^. The burgeoning integration of AI into single-cell discovery has intensified the demand for extensive datasets. For these technologies to perform efficiently, their models must be trained on a comprehensive ensemble of high-quality datasets or fine-tuned on a set of domain-specific biological data^5-8^. Despite the exponential increase in the volume of scRNA-seq data facilitated by the widespread adoption of sequencing technologies, the majority of available data predominantly consists of healthy control samples. This imbalance, exacerbated by economic and ethical considerations, impedes the application of AI in research settings involving rare diseases, specific tissues, and uncommon cell types. Hence, the inevitable need for training datasets significantly burdens research institutions, underscoring the urgency for cost-efficient and time-conscious solutions.

In response, *in silico* data generation strategies, primarily oriented towards global pattern-learning techniques, have been explored along two main avenues: (1) manifold-based and (2) neural network-based approaches. Manifold-based approaches^9-11^ rely on the presumed existence and precise identification of low-dimensional manifolds within scRNA-seq datasets. However, their reliance on dimensionality reduction strategies inevitably comes at the cost of neglecting the nuances of individual cells, thus making them vulnerable to overlooking variations present in high-dimensional data^12^. Additionally, their dependency on the learned manifold confines the generative process to this space, limiting their ability to uncover and explore novel cellular states beyond the existing manifold. More recently, pre-trained generative models such as Generative Adversarial Networks^13^ (GANs) and Variational Autoencoders^14^ (VAEs) from within the AI sphere have ventured to alleviate the data scarcity challenge present in transcriptomic studies (**Fig. 1a-b**). Despite these efforts, they inadvertently enter a paradoxical cycle: their performance remains challenged by the same limited sample availability they seek to overcome^15^. Furthermore, these models are notoriously time-consuming, resource-intensive, and heavily reliant on data-specific tuning of hyperparameters, ranging from architectural features such as the number of layers, layer width, and latent space dimensionality to training configurations, including learning rates, loss functions, and numbers of epochs. These complexities restrict their use to the proof-of-concept stage and make them accessible primarily to advanced users. Hence, this predicament necessitates reevaluating the mounting problem and prompts a pivot towards more foundational yet streamlined solutions.

**Figure 1.**
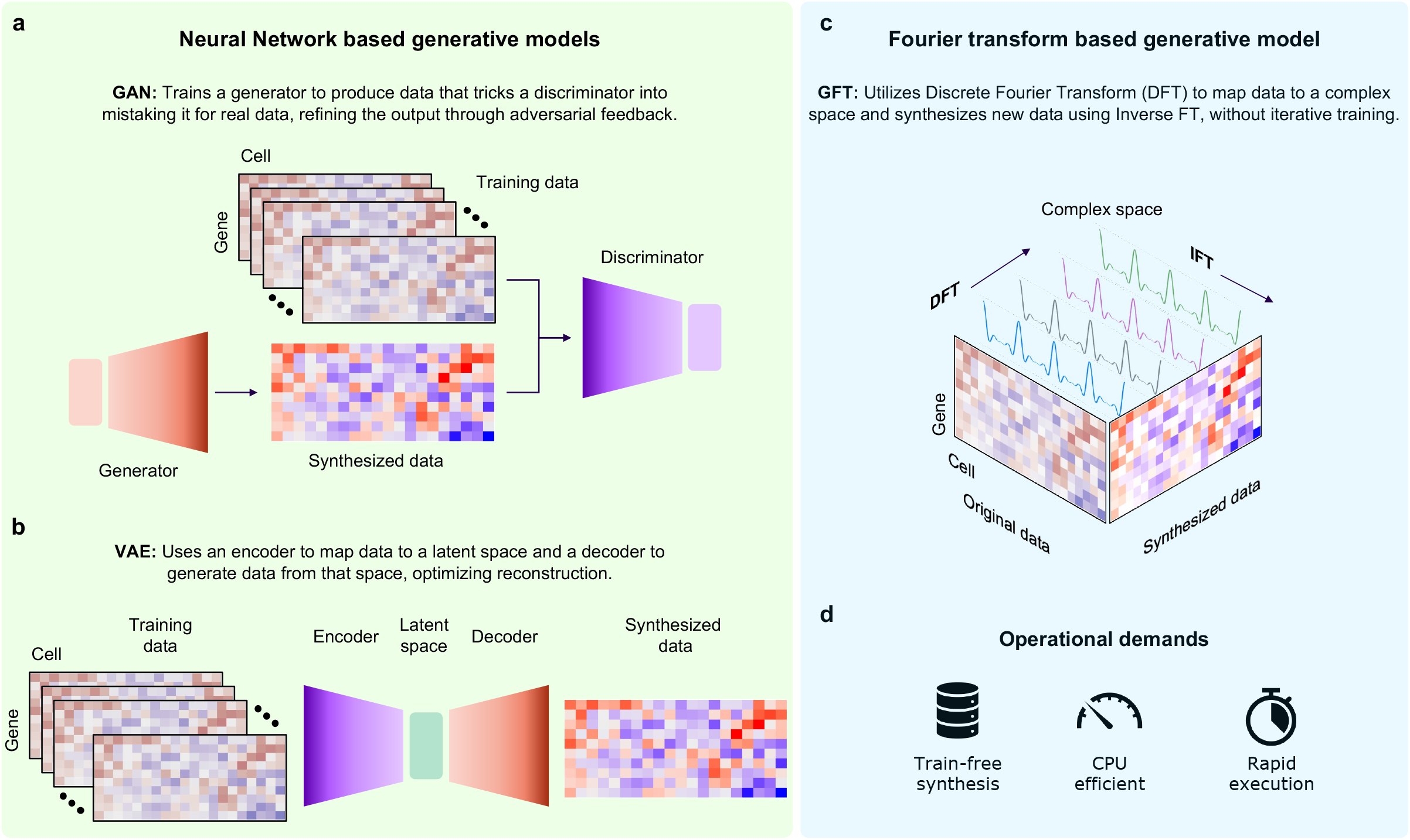
Overview of GAN, VAE, and GFT generative models. **a**, Illustrates the operational frameworks of a Generative Adversarial Network (GAN), focusing on the interactions between the generator and discriminator components for generating synthetic data. **b**, Illustrates the operational frameworks of a Variational Autoencoder (VAE), highlighting the interactions among the encoder, latent space, and decoder for generating synthetic data. **c**, Illustrates the operational framework of a Generative Fourier Transformer (GFT) devised in this study, outlining the transformation of original cell gene expression profiles into a complex space using the Discrete Fourier Transform (DFT) and reconstruction of synthetic profiles using the Inverse Fourier Transform (IFT). **d**, Illustrates the operational resources required by the Fourier transform-based generative model.

We decided to move away from global pattern-learning strategies and instead focus on the cell-centric nature of scRNA-seq data. We posed a fundamental question: Given a cell’s gene expression profile, how can individual cells be synthesized *in silico* with expression profiles that are similar to the original, yet distinct? In response, we aimed to develop an algorithm adept at introducing fine-grained variations into the experimentally observed cell gene expression profiles, thereby fostering the creation of synthetic cell populations that are faithful to the originals. However, these variations cannot be randomly applied to a gene; rather, they must be controlled, robust, and impact the given profile globally to prevent disproportionate alterations in single genes. This endeavor necessitated a mathematical foundation capable of facilitating such transformations.

We were inspired by the versatile application of the Fourier Transform (FT) method across various disciplines^16^. In image processing^17^, the FT is a powerful tool used to enhance and analyze digital images by decomposing them into frequency components. This allows for tasks such as image enhancement, feature extraction, and pattern recognition. In physics^18^, the FT is a powerful tool used to decompose complex physical systems in a given space, such as position, into a more straightforward formulation in another, often momentum space. This transformation simplifies the analysis, for example, of waveforms in quantum mechanics and electromagnetism^19, 20^. Once solutions are obtained, the original physical system can be reconstructed in its original domain using the Inverse Fourier Transform (IFT).

In this context, we mathematically adapted the FT processes to enable the synthesis of single-cell gene expression profiles. We designed scGFT (single-cell Generative Fourier Transformer) framework to host this generative model. The operational conception of scGFT parallels that of VAEs (**Fig. 1b**). Similar to VAEs, which encode input data into a condensed latent space, scGFT utilizes the Discrete Fourier Transform (DFT) to map a cell gene expression profile into a space referred to as “complex space” (**Fig. 1c**). This space comprises a series of complex components (CCs), each capturing unique modes of variation across genes, analogous to the dimensions in a VAE latent space. However, unlike the VAE encoder, the DFT operation is linear and non-reductive, preserving the full information content of the original data.

VAEs generate new data points by first sampling latent variables, then using a decoder to transform these samples back to the original data space - a process governed by the model’s internal parameters, including pre-trained weights and biases (**Fig. 1b**). In a similar vein, scGFT employs the IFT to convert transformed profiles from the complex space back into the gene expression space (**Fig. 1c**). However, unlike the VAE decoder, the IFT operation is linear and reconstructive, accurately restoring the full information content of the original data. Therefore, to ensure the synthesis of unique cellular profiles rather than exact replicas of existing data, the complex space is systematically modified before back-transformation. The IFT then globally propagates the impact of the modified CCs back into the original gene expression profile. Consequently, this process enables the controlled synthesis of an endless array of unique cells, each with a nuanced, deviated profile, while preserving the original cell’s gene expression profile variability.

scGFT offers distinct advantages over conventional generative models in several key aspects. By employing a train-free paradigm, scGFT bypasses the computationally intensive and time-consuming steps associated with training generative models that often require GPU support and extensive cell counts. Unlike neural network-based generative models, which necessitate careful hyperparameter adjustments, scGFT operates on a stable framework that does not rely on data-specific fine-tuning and optimizations. Finally, the FT-based foundation of scGFT eschews the reliance on identifying low-dimensional data manifolds, focusing instead on capturing the intricacies of individual cell expression profiles. Hence, this methodical foundation establishes scGFT as a robust alternative for the *in silico* synthesis of scRNA-seq data, streamlining the data augmentation process and effectively circumventing the common limitations encountered by conventional generative models (**Fig. 1d**).

## Results

A comprehensive and detailed description, including the mathematical principles of the scGFT framework, the implementation steps of cell synthesis, and the analytical methods used in this study, are provided in the Methods section.

### scGFT preserves distinct cellular identities in simulated data during the synthesis process

We aimed to evaluate the quality of cells synthesized by scGFT, focusing on the fidelity of synthesized gene expression profiles to their original counterparts. To achieve this, we employed clustering analysis to rigorously examine entire gene expression profiles, followed by calculating the likelihood that synthesized cells would cluster with their original counterparts. We utilized scRNA-seq simulated datasets to benchmark this criterion, laying a solid groundwork for the analysis.

We generated the simulated data using the Splatter R package^21^, which enabled the creation of three distinct sets of scRNA-seq data, each varying in size and complexity. These included: (1) five datasets, each containing 5,000 cells distributed into five clusters; (2) five datasets, each with 10,000 cells spread across ten clusters; and (3) five datasets, each comprising 15,000 cells divided into fifteen clusters. Cells within each cluster were randomly assigned following a uniform distribution, with each dataset consisting of 10,000 genes. For each of these simulated datasets, we synthesized 50,000 cells to evaluate the scalability and robustness of the scGFT methodology (**Fig. 2a**).

**Figure 2.**
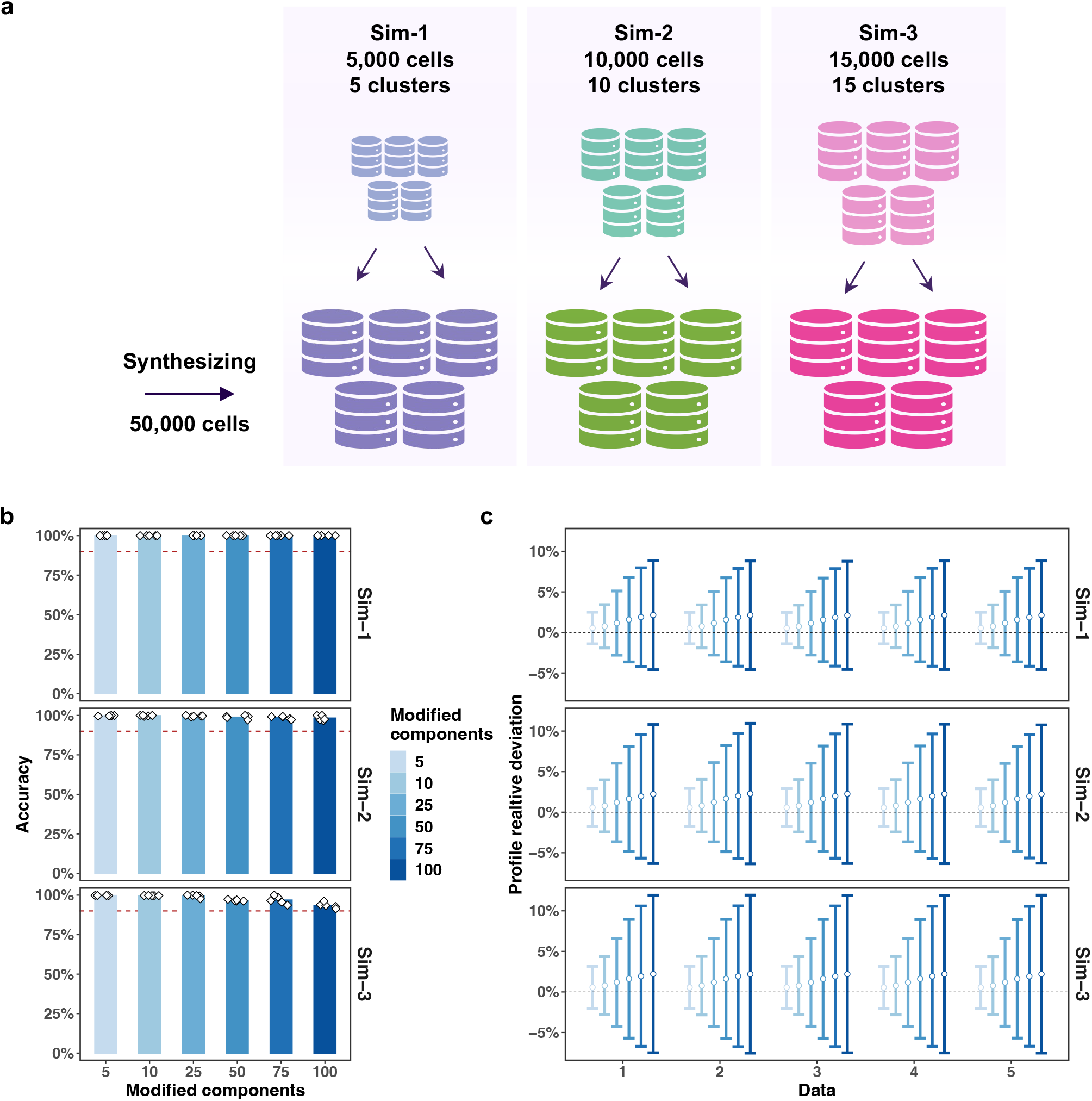
Cells synthesized by scGFT maintain their cellular identities across simulated scRNA-seq datasets. **a**, Illustrates three groups of simulated datasets with varying configurations: (1) Five datasets, each containing 5,000 cells divided into five clusters (sim-1); (2) Five datasets, each with 10,000 cells across ten clusters (sim-2); and (3) Five datasets, each comprising 15,000 cells sorted into fifteen clusters (sim-3). Each cluster within these datasets was populated through random assignment following a uniform distribution, with each dataset featuring 10,000 genes. For each dataset, 50,000 cells were synthesized with varying numbers of complex components affected (5, 10, 25, 50, 75, and 100) using scGFT. **b**, Bar plots depicting the proportion of synthesized cells that accurately clustered with their corresponding original cells, for each dataset (diamonds). The x-axis represents the number of complex components modified, and the y-axis shows the clustering accuracy. Each row corresponds to one of the three groups of datasets. A horizontal dashed red line highlights the 90% accuracy threshold. **c**, Error bars (mean ± standard deviation) depicting the impact of modifying varying numbers of complex components (color-coded) on the deviation of gene expression profiles in synthesized cells from their original counterparts (y-axis), for different datasets (x-axis). Each row corresponds to one of the three groups of datasets.

Synthesis was performed (n=50,000 cells) for each dataset, with varying numbers of CCs (5, 10, 25, 50, 75, and 100) modified to examine the impact of different levels of modification on synthesis fidelity. Synthesized cells were then integrated with the original data and re-clustered. We assessed the quality of the synthesis process by calculating the proportion of synthesized cells that correctly clustered with their corresponding original cells. This evaluation revealed a high accuracy, exceeding 92% in each dataset and averaging above 97% across all numbers of modified components and simulated data (**Fig. 2b**).

We noted a decrease in accuracy in our analysis of the largest simulated dataset as the number of modified CCs increased. To investigate the impact of altering the number of modified CCs, we sought to quantify the divergence of gene expression profiles between synthesized cells and their originals. We calculated the relative average deviation of gene expression between synthesized and original cells for varying numbers of modified CCs (5, 10, 25, 50, 75, and 100) in the simulated datasets. The results demonstrated an anticipated increase in deviation magnitude as more CCs were modified, aligning with the theoretical underpinnings of the scGFT method (**Fig. 2c**). This analysis elucidates the observed decrease in clustering performance, which stems from the increased disparity between synthesized cells and their originals due to the greater number of modified CCs. Moreover, the analysis revealed a slight positive shift in the deviation distributions (**Fig. 2c**). This shift is attributed to the Rectified Linear Unit (ReLU) function, implemented to clip negative values introduced during synthesis to zero.

Overall, this analysis confirms the controlled basis of the variations introduced by scGFT and the integrity of the synthesized cells’ expression profiles by maintaining their distinct cellular profiles in proximity to their original counterparts.

### scGFT augment experimental scRNA-seq data maintaining intrinsic cellular characteristics

To assess the performance of scGFT in practical applications, we extended our investigation to include experimental scRNA-seq data. We utilized the dataset PRJEB44878^22^, which comprises 34,200 processed cells derived from primary small airway epithelial cells (SAECs) from both healthy individuals (n=3) and patients with chronic obstructive pulmonary disease (COPD) (n=3). These SAECs were subjected to in vitro expansion and differentiation into pseudostratified epithelia via air-liquid interface (ALI) conditions. To model smoke-induced injuries in the small airways of healthy non-smokers and COPD smokers, the fully differentiated SAEC ALI cultures underwent exposure to either whole cigarette smoke over a period of four consecutive days or to ambient air serving as the control.

For this analysis, scGFT synthesized new cells at varying expansion coefficients - single (1x), double (2x), and triple (3x) the number of original cells - through modification of 5, 10, 25, 50, 75, and 100 CCs. Utilizing UMAP for a qualitative evaluation, we projected synthesized and real cells onto the embedded manifold. This visualization demonstrated a substantial overlap across all expansion coefficients (**Fig. 3a**; synthesized data with 100 CCs modified). Additionally, this illustration indicates that the synthesis process did not introduce any non-biological artifacts. This evaluation was further quantitatively corroborated by clustering analysis, in which both synthesized, and original, cells demonstrated an alignment exceeding 92% accuracy, with an average above 94% across all numbers of modified components and expansion coefficients (**Fig. 3c**). We note that to process the experimental scRNA-seq data, we adhered to the same analysis procedures both before and after synthesis, following the standard Seurat pipeline^23^.

**Figure 3.**
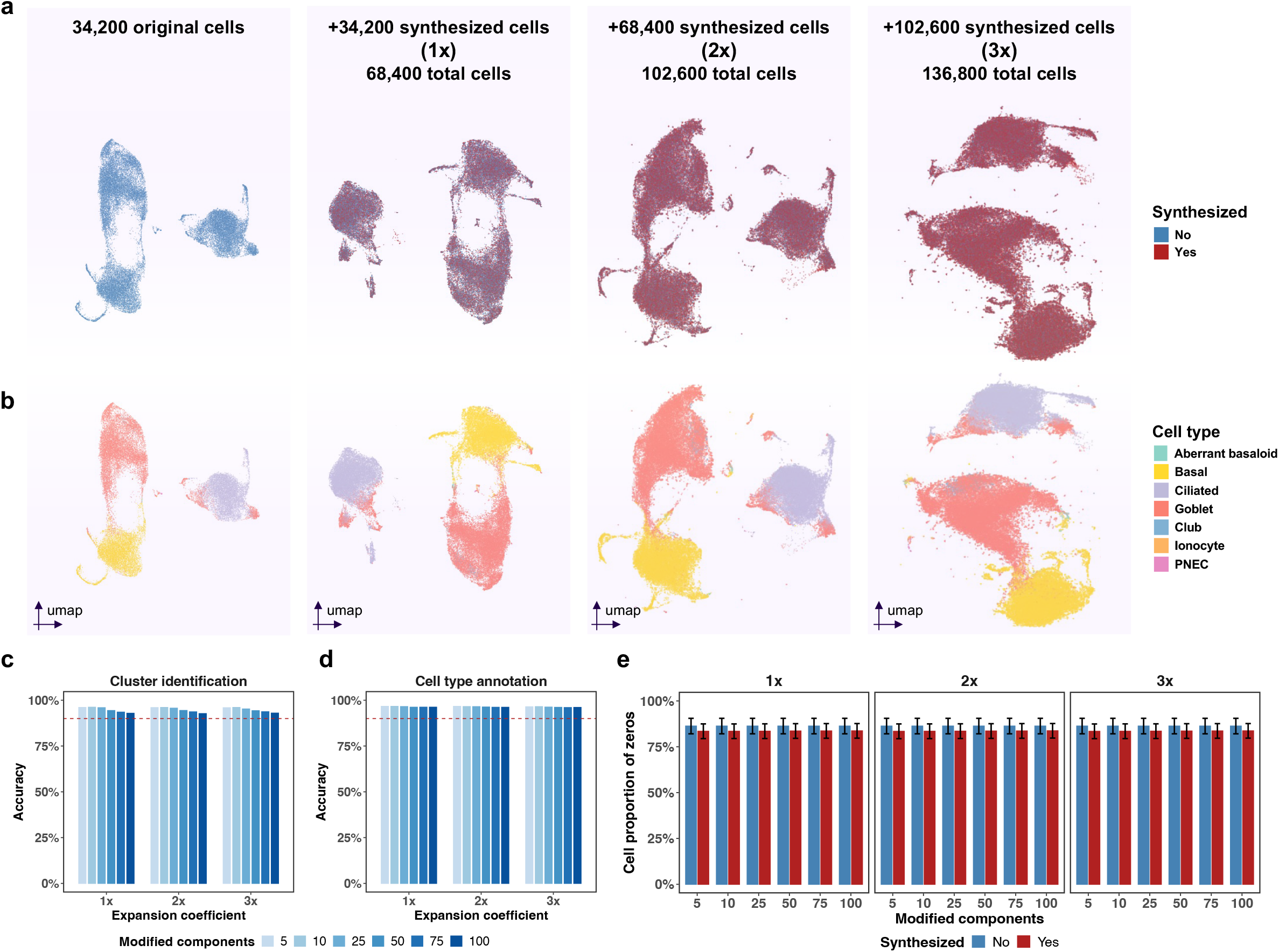
Cells synthesized by scGFT maintain their cellular characteristics across experimental scRNA-seq datasets. **a**, UMAP representation of the PRJEB44878 dataset comprised of 34,200 processed single cells from primary small airway epithelial cells. The first column shows the original dataset. Subsequent columns illustrate original cells (blue) combined with cells synthesized by scGFT (red), modifying 100 complex components under different expansion coefficients: 1x synthesis resulting in 68,400 total cells (second column), 2x synthesis resulting in 102,600 total cells (third column), and 3x synthesis resulting in 136,800 total cells (fourth column). **b**, UMAP visualization of the same dataset across different expansion coefficients, color-coded according to annotated cell types. **c**, Bar plots depicting the proportion of synthesized cells that accurately clustered with their corresponding original cells, shown for each expansion coefficient (x-axis). Each color denotes the number of complex components modified, and the y-axis measures clustering accuracy. A horizontal dashed red line highlights the 90% accuracy threshold. **d**, Bar plots depicting the proportion of synthesized cells correctly annotated as identical to their corresponding original cell types, presented for each expansion coefficient (x-axis). Each color indicates the number of complex components modified, and the y-axis measures annotation accuracy. A horizontal dashed red line indicates the 90% accuracy threshold. **e**, Bar plots showing cell data sparsity with the proportion of zeros calculated for original (blue bars) and synthesized (red bars) cells across various numbers of complex components modified (x-axis) and different expansion coefficients (1x, 2x, and 3x, represented in left, middle, and right panels, respectively). Error bars show the mean ± standard deviation of the proportions calculated for each category.

Genes with high variability manifest biological variations indicative of distinct cellular states. We sought to further substantiate the preservation of inherent variability in the original cells within the synthesized data. We calculated the proportional overlap of the top 2,000 highly variable genes identified from the original and synthesized data across six different component modifications (5, 10, 25, 50, 75, and 100 components) and three expansion coefficients (1x, 2x, and 3x). This assessment successfully demonstrated a high average overlap of 94 ± 0.7% (mean ± standard deviation) between two gene sets.

Cell type annotation is a crucial downstream analysis in single-cell RNA sequencing, offering insights into the cellular heterogeneity within samples. We extended our analysis to evaluate the consistency of cell types in synthesized cells relative to the originals. To ensure an unbiased assessment, we utilized Sargent^24^, an automated, cluster-free, score-based annotation method that classifies cell types based on distinct gene expression markers. We curated a comprehensive list of canonical epithelial subtype markers by leveraging experts’ knowledge and the literature^22, 25^ (**Supplementary File 1**). Sargent identified distinct epithelial cell subtypes, including goblet cells (n=15,472), ciliated cells (n=9,515), and basal cells (n=9,035), as well as less abundant subtypes such as club cells (n=94), aberrant basaloid cells (n=47), pulmonary neuroendocrine cells (PNECs) (n=21), and ionocytes (n=16) (**Fig. S1**). The UMAP visualizations again demonstrated a strong overlap between annotated synthesized cells and the original dataset across all expansion coefficients (**Fig. 3b**: synthesized data with 100 CCs modified). This overlap was quantitatively supported by cell type re-annotation analysis, where both synthesized and original cells demonstrated concordance exceeding 95% accuracy in cell type labeling, with an average above 96% across all numbers of modified components and expansion coefficients (**Fig. 3d**).

Data sparsity, represented by zeros, is a characteristic feature of single-cell RNA sequencing datasets. To evaluate this aspect of the synthesis process, we compared the number of zeros in synthesized and original cells across various numbers of modified components and expansion coefficients. This analysis revealed that the synthesized data exhibited a sparsity level slightly lower than the original data (85% versus 86%, respectively). The trend remained similar across all numbers of modified components and expansion coefficients, affirming that scGFT maintains the integrity of gene expression data without artificially denoising or extensively (< 1%) imputing (**Fig 3e**). Overall, the findings from experimental scRNA-seq datasets highlight the scGFT capability to effectively augment scRNA-seq data in practical settings while preserving intrinsic cellular characteristics and biological plausibility, both critical elements for robust downstream analytical applications.

### scGFT synthesizes distinct cell populations solely from individual cell gene expression profiles

Analyzing rare cell types in scRNA-seq data presents a formidable challenge due to their low abundances, significantly impacting statistical power and analytical precision. Addressing this challenge touches on the core purpose of this study, which concerns the ultimate capacity of generative models to synthetically generate biologically plausible cellular populations solely from the gene expression profile of an individual cell.

We sought to assess the effectiveness of scGFT in addressing this challenge by focusing on rare epithelial subtypes in the PRJEB44878 experimental data, including *SCGB3A2*^+^ club cells, *MMP7*^+^ aberrant basaloid cells, GRP^+^ PNECs, and *FOXI1*^+^ ionocytes, each comprising less than 0.3% of the population. An individual cell from each population was randomly selected for analysis. Subsequently, we modified 100 CCs and synthesized 5,000 cells to evaluate scGFT’s ability to expand these rare cell profiles. This synthesis process effectively generated discrete cell populations, each remaining separate from others (**Fig. 4a-d**). Quantitative support for this augmentation process came from annotation analyses of the integrated data, where both synthesized and original cells demonstrated a high degree of agreement in cell type labeling, with accuracy exceeding 98% on average across the four examined cell types.

**Figure 4.**
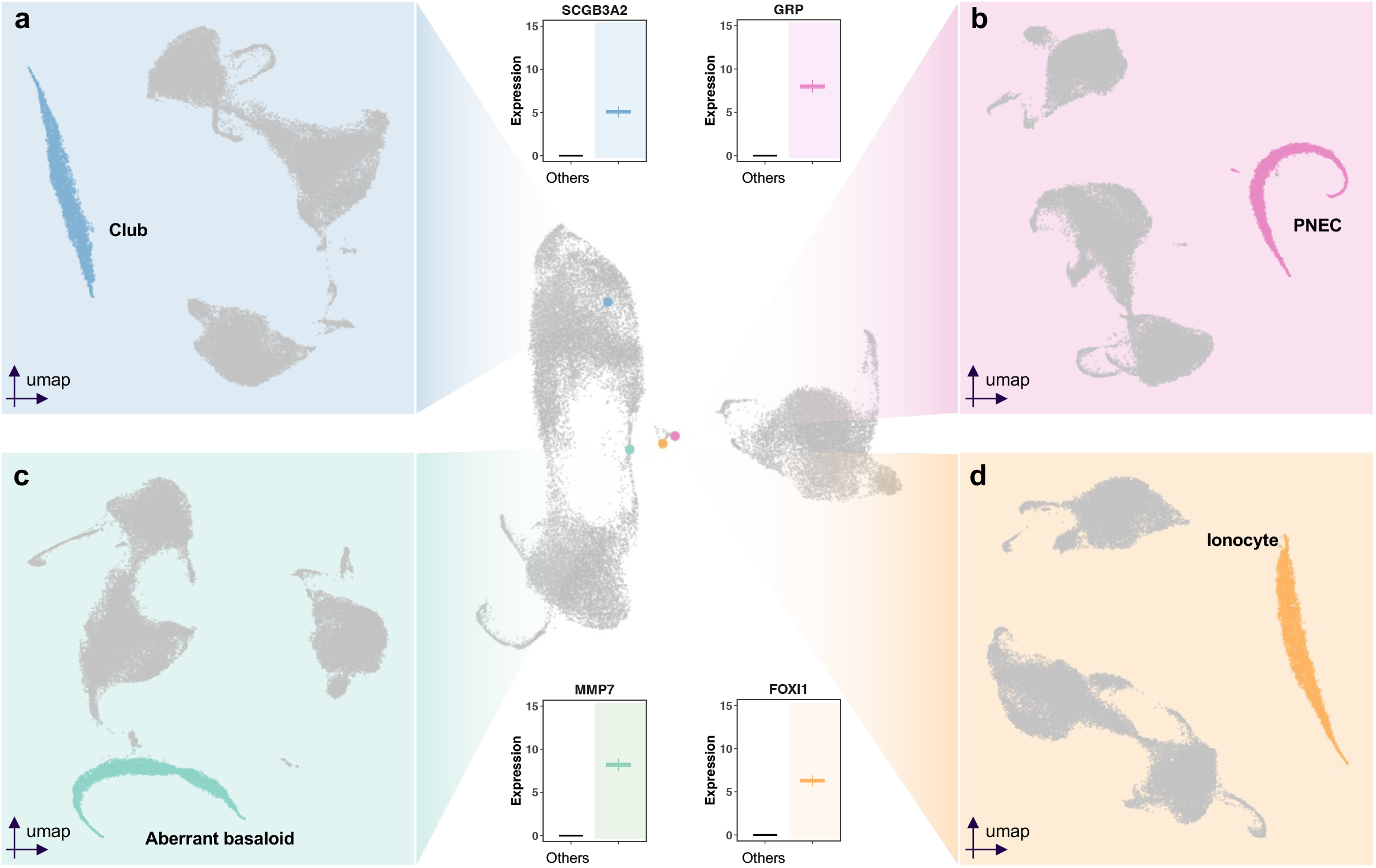
scGFT augments rare cell types by synthesizing cellular populations from individual cell profiles. UMAP representation of the PRJEB44878 dataset (center panel) with randomly selected individual rare cell types highlighted: a club cell (blue), a pulmonary neuroendocrine cell (PNEC; pink), an aberrant basaloid cell (green), and an ionocyte (orange). Modifying 100 complex components, 5,000 cells synthesized by scGFT for each of the selected cell types are represented by UMAPs for club cells, PNECs, aberrant basaloid cells, and ionocytes depicted in **a-d**, respectively. Cells from other types are colored grey to distinguish the synthesized populations. The box plots show the expression levels of the selected markers for each cell type. The upper and lower bounds of the boxplot represent the 75th and 25th percentiles, respectively. The center bars indicate the medians, and the whiskers denote values up to 1.5 interquartile ranges above the 75th or below the 25th percentiles.

Hence, this analysis unveils the unique ability of scGFT to synthesize a distinctive population of cells exclusively based on the expression profile of an individual cell, highlighting a significant leap forward and surpassing the capabilities of current state-of-the-art generative methods.

### scGFT enhances network analyses for identifying gene programs in human alveolar epithelium

One of the primary goals in transcriptomics studies is to identify sets of functionally coherent genes and their associations with various biological contexts (e.g., cell states, tissues, treatments, and disease states), thereby providing insights into the pathways that govern cellular processes. Network inference strategies are employed to elucidate these gene associations and interactions. However, the effectiveness of these strategies is often compromised in rare cell types due to insufficient observations, which limit the comprehensive modeling of gene-gene interactions.

To illustrate the application of scGFT in enhancing detailed and statistically robust network inference analyses, we utilized the dataset GSE178360. This dataset consists of 7,160 processed cells obtained from microdissected healthy lung distal airways. For consistency, we retained the original cell type annotations provided by the authors^26^. Our objective was to perform a data-driven identification of cell type-specific pathways in rare epithelial alveolar types, namely AT0, AT1, and AT2. Consequently, we aimed to infer gene-gene association networks for *SCGB3A2*^+^*SFTPC*^+^ AT0, *SCGB3A2*^-^*SFTPC*^-^ AT1, and *SCGB3A2*^-^*SFTPC*^+^ AT2. These subtypes each constitute less than 5% of the population (n=287, 275, and 327 cells, respectively), underscoring the challenge of deriving informative networks due to limited observations.

Using scGFT, we synthesized 1,000 cells for each AT subtype by modifying 10 CCs. We employed UMAP for qualitative evaluation and compared the expression profiles of selected AT marker genes, demonstrating the fidelity of the synthesized profiles to their originals (**Fig. 5a-b**). This evaluation was further quantitatively corroborated by comparing the differentially expressed genes (DEGs) among subtypes. We conducted differential expression analysis (DEA) using AT0 as the baseline, establishing thresholds of adjusted p-values below 0.05 and log2 fold changes exceeding 1. This analysis resulted in a 96% overlap in DEGs for AT1s (507 DEGs in original and 560 DEGs in synthesized cells) and an 85% overlap in DEGs for AT2s (60 DEGs in original and 68 DEGs in synthesized cells). The overlap was calculated as the intersection of two gene sets divided by the average of the lengths of each gene set.

**Figure 5.**
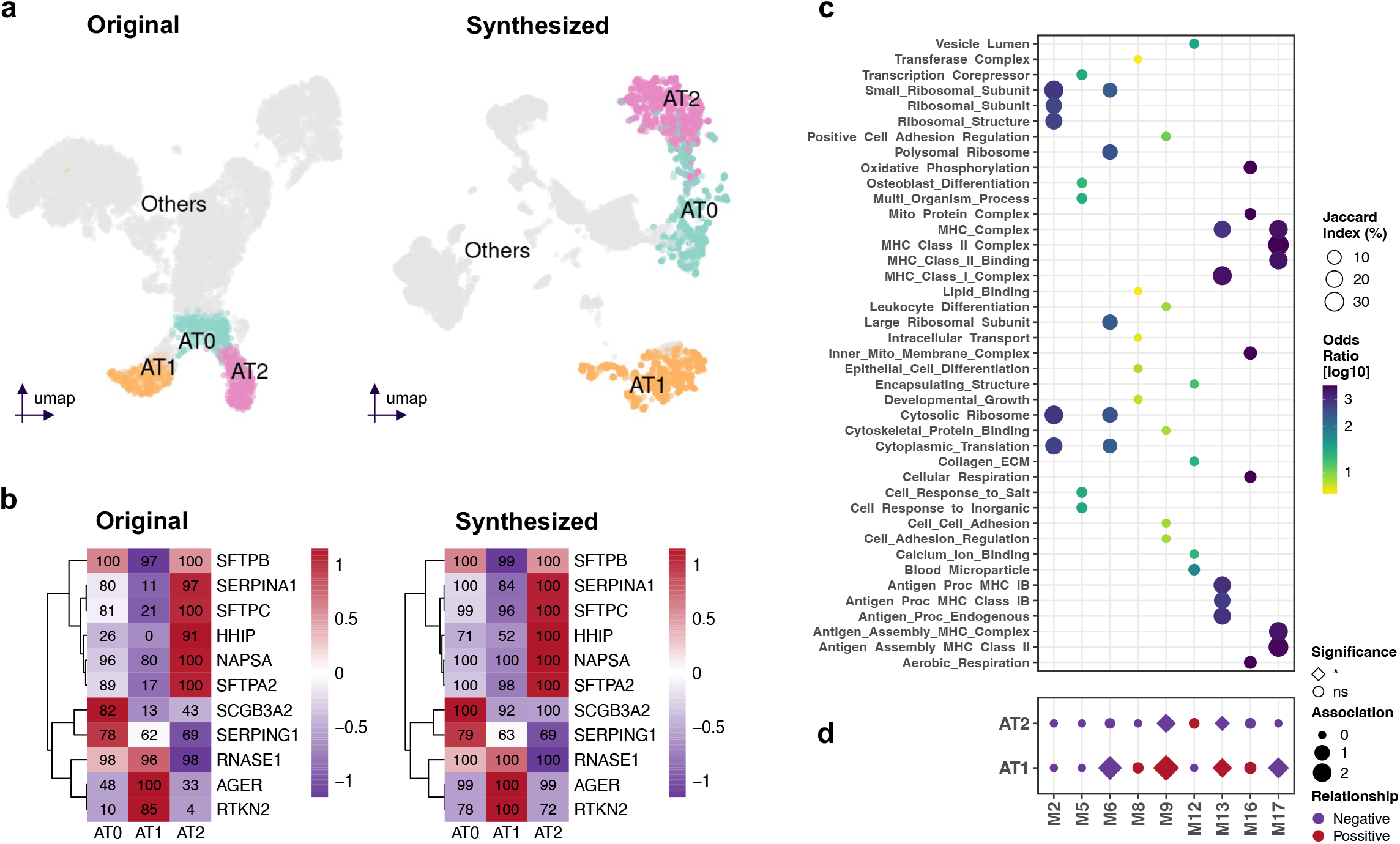
scGFT facilitates the identification of gene programs through enhanced network analyses in human alveolar epithelium. (**a**) UMAP representation of the GSE178360 dataset comprised of 7,160 processed cells obtained from microdissected healthy lung distal airways. The original UMAP is shown in the left panel, while the right panel illustrates the combined original and 1,000 synthesized cells for each alveolar subtype: AT0, AT1, and AT2. Cells from other types are colored grey. (**b**) Heatmap displaying the average expression level of selected marker genes for each of the different alveolar subtypes in the GSE178360 dataset. The value in each element indicates the percentage of cells expressing the given gene in each of the different alveolar subtypes. (**c**) Gene enrichment analysis for annotating inferred module functions using the Gene Ontology (GO) database. The x-axis corresponds to the selected inferred module identifiers, while the y-axis represents the identified GO terms. The color bar in the legend indicates the odds ratio, representing the strength of association between the modules and GO terms. The circle sizes in the legend display the Jaccard index, measuring the similarity between two lists of genes. Fisher’s exact test was employed to determine statistical significance. The results are presented for the top five GO modules with the highest odds ratios and an adjusted p-value below 0.05. (**d**) Quantification of gene module associations with biological states. A positive coefficient implies that higher expression of a module is associated with the state under study (AT1 and AT2), while a negative coefficient implies an association with the baseline state (AT0). The sizes of the data points indicate the magnitude of the association. Significantly (\**p*-value<0.05) associated modules are shown as diamonds, while non-significant (ns) modules are shown as circles.

Next, we inferred gene-gene association networks for the combined original and synthesized data for each AT subtype, resulting in graphs containing 5,879 genes for AT0, 4,551 genes for AT1, and 6,936 genes for AT2. We then identified modules as clusters of genes exhibiting dense interconnections within these graphs, yielding 24 modules in AT0, 28 modules in AT1, and 16 modules in AT2. These modules were subsequently merged into 18 unified modules (M1-M18) (**Supplementary File 2**), distinguishing between general modules shared across all ATs and those specific to individual subtypes.

We then conducted gene enrichment analysis to annotate the function of these modules, utilizing gene ontology (GO) terms (**Supplementary File 2**). The analysis revealed substantial overlap between the inferred modules and pathways essential for the respiratory function (**Fig. 5c**). Pathways consisting of ribosomal proteins were found in M2 and M6, an expected and reassuring finding given that such modules are fundamental to basic cellular functionality. Genes associated with cellular responses to inorganic substances were found in M5, highlighting the importance of these responses for lung tissues frequently exposed to such substances. Epithelial cell differentiation genes, critical for the state transitions of ATs, were identified in module M8. Intriguingly, the expression of this module is positively associated with AT0 and AT1, and negatively correlated with AT2 (**Fig. 5d**). This finding aligns with the original GSE178360 paper’s claim that AT2 cells differentiate into AT0 and AT1 cells, suggesting a downregulation of epithelial differentiation genes in AT2 as they transition to a more differentiated state in AT0 and AT1. Cell-cell adhesion relevant genes were found in M9, playing a crucial role in forming tight junctions and maintaining the integrity of the air-blood barrier in alveolar subtypes. Collagen and extracellular matrix components, vital for the structural integrity of lung tissue including alveoli, were found in M12. Pathways involved in antigen processing, including major histocompatibility complex (HLA) class I and II genes, were identified in M13 and M17, respectively, with M13 positively and M17 negatively associated with AT1 (**Fig. 5d**). Lastly, genes involved in aerobic respiration and adenosine triphosphate production, essential for gas exchange in lung cells, particularly alveolar cells, were identified in M16.

Overall, this analysis demonstrates the potential scGFT holds to facilitate detailed and statistically robust downstream analysis of scRNA-seq data.

### scGFT computational efficiency obviates the requirement for high-performance computing

Computational efficiency is a critical and costly attribute in research settings. We evaluated the computational performance of the scGFT by recording the time it took to synthesize cells from both simulated and experimental datasets. These tests were conducted on a single-core setup with a 2.9 GHz processor and 64 GB of memory.

In the analysis of simulated data, where the target number of synthesized cells was fixed at 50,000, we observed that the synthesis time required by the scGFT model varied based on the size of the original data and the number of modified CCs. Specifically, for the smallest dataset containing 5,000 cells, approximately 0.8 minutes were required to synthesize 50,000 cells when only five components were modified. This duration increased to approximately one minute when 100 components were modified. Similar patterns were observed for larger datasets containing 10,000 and 15,000 cells, with the longest runtime of approximately 1.5 minutes for maximum component modifications (n=100) (**Fig. 6a**). The minimal increase in runtime in the scenario with a fixed number of CC modifications and a fixed number of synthesized cells is attributed to the DFT process applied to an increasing number of original cells.

**Figure 6.**
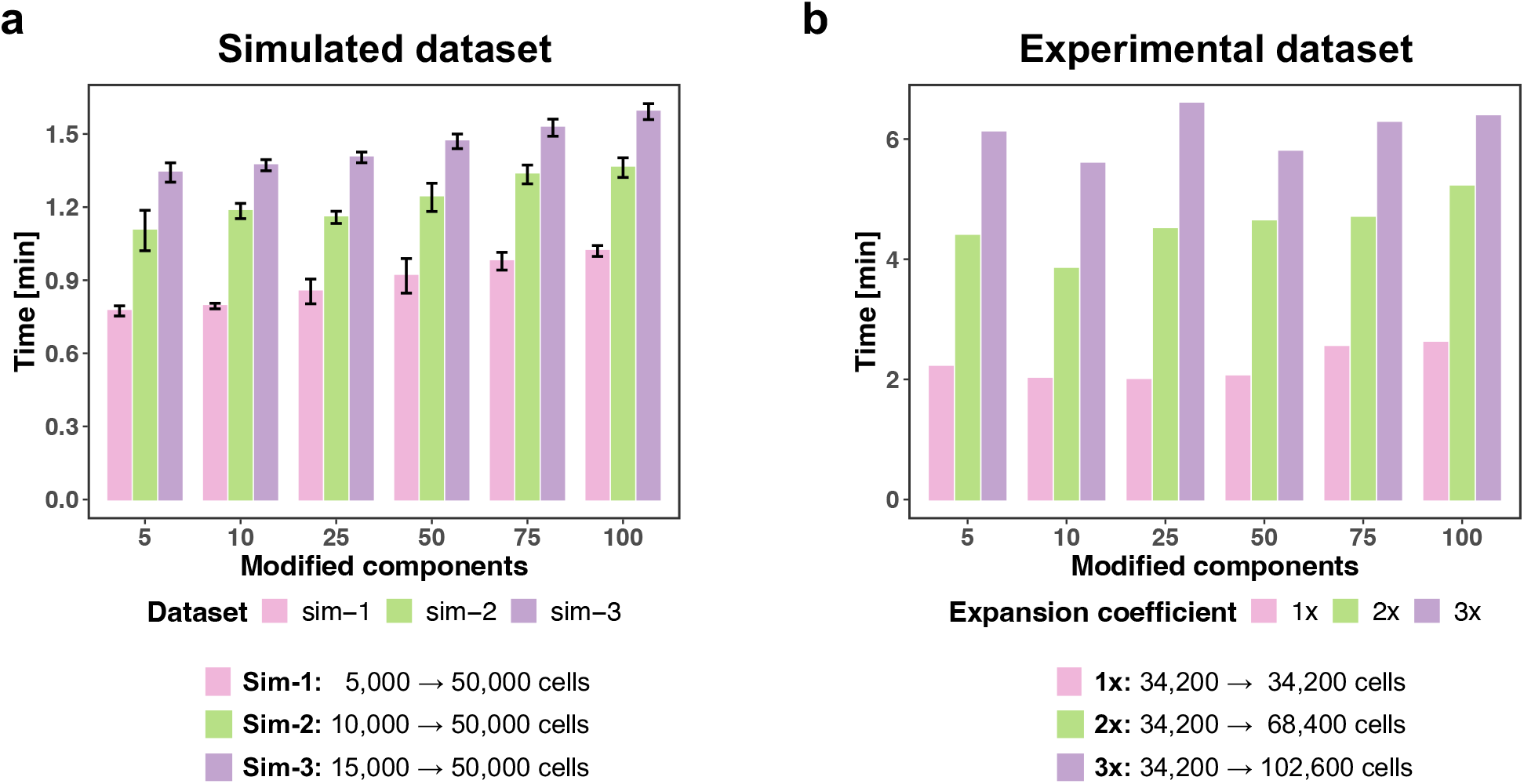
scGFT efficiently synthesizes large numbers of cell gene expression profiles. Runtime (y-axis) of the synthesis process, executed on a uniprocessor system (equipped with a 2.9 GHz processor and 64 GB of memory), is plotted against the number of complex components modified (x-axis) in simulated datasets (**a**) and PRJEB44878 experimental dataset (**b**). The color legend in (**a**) indicates different simulated data groups, and in (**b**), it denotes the expansion coefficients of the synthesis process. Error bars (mean ± standard deviation) in (**a**) are calculated across datasets (n=5) in each simulated data group.

In the case of experimental data, the computational time was influenced by the number of cells to be synthesized and the number of components modified. The synthesis time ranged from approximately two minutes for a single expansion coefficient (1x; 34,200 cells) with minimal component modifications (n=5), to nearly six minutes for the highest expansion coefficient (3x; 102,600 cells) and maximum component modifications (n=100) (**Fig. 6b**).

This analysis reveals that the runtime of the scGFT algorithm scales additively with the number of original cells (*o*) and multiplicatively with both the number of synthesized cells (*σ*) and the number of modified CCs (*κ*). Consequently, the computational complexity is represented as *O*(*o* + *σk*). Additionally, this analysis demonstrates the algorithm’s efficiency on uniprocessor systems without requiring GPU acceleration.

## Discussion

As the adoption of AI-based technologies in research settings continues to expand, the demand for extensive datasets to effectively train these models presents a persistent challenge. This limitation hinders the predictive power of AI, especially in single-cell transcriptomics, where high-quality training datasets are often scarce, such as in the study of uncommon cell types, specific tissues, or rare diseases. In this study, we present the scGFT framework, which emerges at the intersection of AI and scRNA-seq data, directly addressing data scarcity by generating realistic synthetic single cells.

scGFT comprises a streamlined computational framework grounded in mathematical rigor, leveraging the principles of the Fourier Transform. This rigor enables a successful in silico synthesis process, carefully tailoring the original gene expression profile while preserving the variability of the original population. We validated the performance of scGFT through comprehensive assessments using both simulated and experimental data across various scenarios. This validation illustrates scGFT’s capability to generate synthetic cells that preserve the intrinsic characteristics delineated in the original scRNA-seq data, as evidenced by thorough evaluations, including clustering and cell-type analyses. Furthermore, leveraging experimental data, we probed established features of scRNA-seq data, confirming that scGFT generates sparsity rates similar to those observed in real experimental data, does not denoise or impute gene expression information, and effectively discards potential non-biological artifacts during the synthesis process.

scGFT facilitates the synthesis of an unlimited array of unique cells from experimentally observed cell gene expression profiles. This stochastic process stems from three mechanisms: (1) the random selection of cells for synthesis, (2) the random selection of CCs for modification, and (3) the random sampling of modification factors. In addition, scGFT obviates the need for manifold reduction techniques, extensive training phases, and complex hyperparameter tuning strategies common in conventional generative models. These features ensure scGFT swift execution without requiring high-performance GPU resources, making it accessible across various computing settings.

scGFT enables controlled synthesis through the modification of CCs. An increase in the number of modified components amplifies the profile disparity between synthesized cells and their originals, potentially steering them toward previously uncharted gene expression profiles. While this exploration is integral to the premise of generative models, which aim to uncover cellular states absent from existing datasets, it requires rigorous investigation to ensure the biological relevance of these novel states. It should also be noted that the post-synthesis smoothing mechanism implemented in scGFT attenuates excessive deviations and potential artifacts, ensuring that the modifications remain within a biologically relevant range.

scGFT is capable of initiating the synthesis process from an individual single cell. This feature surpasses the capabilities of current state-of-the-art generative models, which require a substantial amount of data for training. In addition, scGFT holds the potential to support the development of newly emerged foundational models for scRNA-seq, particularly in scenarios where training or fine-tuning is challenging due to scarce sample availability. Integrating large-volume synthesized datasets for training can mitigate potential artifacts that arise when training exclusively with scarce real datasets, which often has inconsistencies from different laboratories, such as contamination and high variance in sequencing read depth. This strategy enables models to learn uniform and meaningful representations of cells and genes.

scGFT presents the feasibility to generate anonymized, disease-specific synthetic single-cell patient repositories, thereby addressing the financial and ethical constraints often encountered in clinical settings. This capability supports the development of robustly trained AI-based classifiers, e.g. for patient stratification, which are essential for designing effective clinical trials. Consequently, scGFT serves as an ethically sound and cost-effective catalyst for advancements in therapeutic discovery, enabling more efficient and precise medical research and development.

The methodology underlying scGFT is versatile and can be applied to any count matrix data, including those derived from various omics technologies, such as bulk RNA-seq data. This adaptability extends to medical imaging research, where data typically consists of pixel intensity matrices. Potentially, by employing a 2-dimensional DFT (2D-DFT) on the grid of pixel intensities, the image is converted from the spatial domain to the frequency domain, capturing frequency components in both row-wise and column-wise directions. Parallel to the scGFT concept, systematically modifying a subset of these frequency components and subsequently applying the 2D-IFT allows for the synthesis of new images with controlled variations. This approach enables researchers to generate realistic synthetic medical images, addressing data scarcity and enhancing the training of AI-based models^27^ in medical imaging applications.

In conclusion, designed to be compatible with the widely-used Seurat framework, the scGFT framework not only enhances its integration into existing bioinformatics pipelines but also contributes to the democratization of scRNA-seq data, representing a transformative force in precision medicine.

## Methods

### Discrete and inverse Fourier transform of a gene expression profile

The gene expression profile is quantified for each cell as a discrete series, *x*[*n*], with *n* indexing genes (0 to *N* − 1, *N* being the total genes). The Discrete Fourier Transform (DFT) is applied to transform each cell gene expression profile to the complex space:

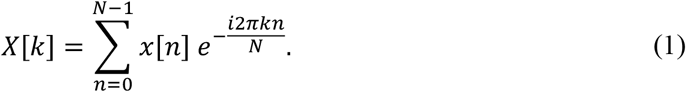

Here, *X*[*k*] denotes the complex value *a* + *ib*, encapsulating the amplitude, *a*, and phase, *b*, for complex component *k*, with *i* as the imaginary unit. The Inverse Fourier Transform (IFT) is then utilized to revert the complex space data, *X*[*k*], back to the original gene expression profile *x*[*n*]:

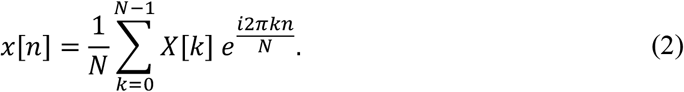

This operation satisfies both additivity and homogeneity principles, and therefore is linear. Additionally, it preserves the order and magnitude of the input gene expression profile.

### IFT propagates complex space modifications globally to gene profiles

We investigate the implications of amplitude adjustments to specific complex components (CCs) within a FT series and their subsequent effects on gene expression profile reconstruction.

#### Lemma

Modification of the amplitude of a specific complex component, *X*[*m*], within a Fourier-transformed series systematically impacts the entire reconstructed gene expression profile.

**Proof:** The modification of the *m*-th complex component, *X*[*m*], of a Fourier-transformed series by a multiplicative factor *A* yields the modified component *AX*[*m*]. The impact of this alteration on the reconstructed profile, *x*’[*n*], as manifested through IFT can be expressed as:

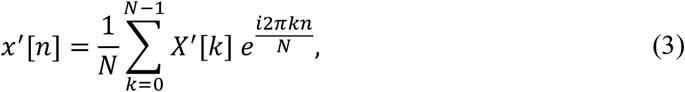

where, *X*’[*k*] denotes the modified CC spectrum. For all *k* except for *m, X*’[*k*] = *X*[*k*]; however, for *m, X*’[*m*] = *AX*[*m*]. The reconstructed signal, *x*’[*n*], can thus be expressed as the sum of the original signal (except the *m*-th component) plus the contribution from the modification:

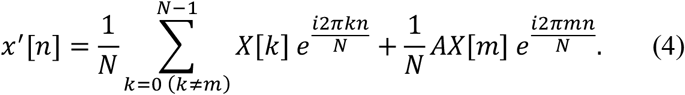

By adding and subtracting the unmodified *m*-th component:

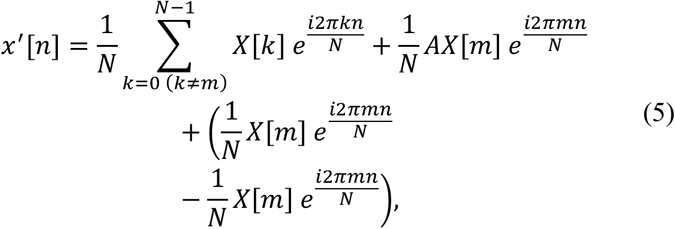

the expression can be simplified in the form of:

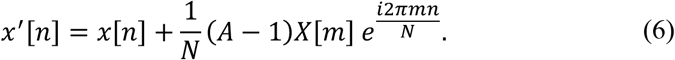

This formulation elucidates that modifying the complex component *X*[*m*] introduces an additional term to the reconstructed signal *x’*[*n*], which is dependent on *n*, the gene index in the original domain. Thus, the term (*A* − 1)*X*[*m*]*e*^*i*2*πmn/N*^ encapsulates the impact induced by the alteration of *X*[*m*], influencing all values of *n*. This feature is pivotal, demonstrating the systematic propagation of a modification across the entire measured transcriptomic landscape on a mathematically rigorous basis.

### The conjugate symmetry principle of FT preserves the real-valued nature of gene expression profiles

Conjugate symmetry is a fundamental principle in the application of FT technique to real-valued datasets. Therefore, when modifying the amplitude of a CC, it is crucial to apply the same modification to its conjugate pair to maintain the real-valued nature of the gene expression profile upon performing the IFT. The following proofs substantiate this critical characteristic.

#### Lemma

Given that *x*[*n*] is real, the Fourier Transform *X*[*k*] of *x*[*n*] exhibits a specific symmetry, known as conjugate symmetry, defined by:

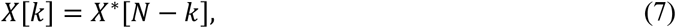

for all *k*, where *X*^*^[*N* − *k*] is the complex conjugate of *X*[*k*]. The asterisk (^*^) denotes the complex conjugate of a complex number, where the real parts are identical, and the imaginary parts are negatives of each other.

**Proof:** The discrete Fourier transform of a sequence *x*[*n*], where *n* = 0, 1, …, *N* − 1 and *N* represents the total number of genes, is given by:

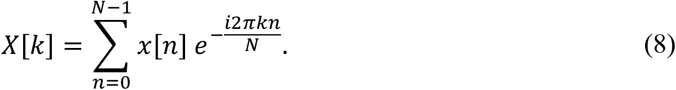

The component *N* − *k* can be expressed as:

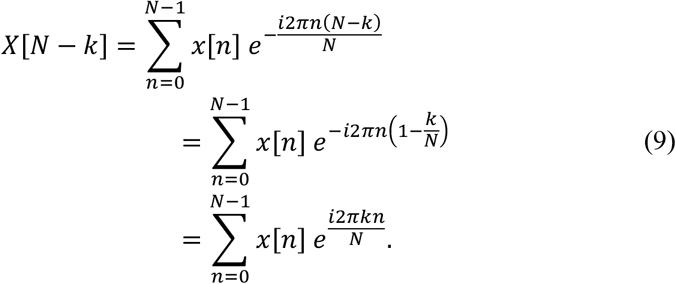

This equality holds because *e*^-*i*2*πn*^ = 1 (as *e*^-*i*2*πn*^ represents a full rotation in the complex plane). Taking complex conjugate of both sides:

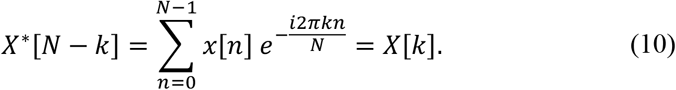

Here, given the real-valued nature of the gene expression profile, *x*^*^[*n*] = *x*[*n*].

#### Lemma

The conjugate symmetry ensures that performing the IFT results in a real value, adhering to the physical nature of the data represents.

**Proof:** Given the DFT of a real-valued sequence *x*[*n*], denoted by *X*[*k*], its IFT is represented as:

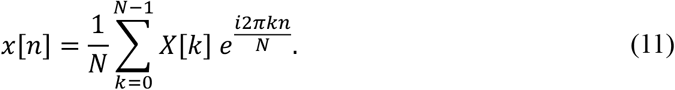

Applying Euler’s formula, *e*^*i*^θ = cos(θ) + *i* sin(θ), the contributions of a generic *X*[*k*] within the summation is:

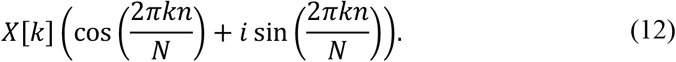

Subsequently, the contribution of its conjugate *X*[*N* − *k*] to the sum, utilizing the conjugate symmetry property, takes the form:

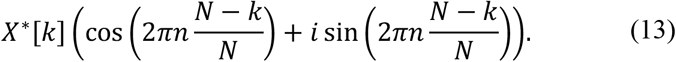

Given that *X*^*^[*k*] is the complex conjugate of *X*[*k*], if *X*[*k*] = *a* + *ib*, then *X*^*^[*k*] = *a* − *ib*. Therefore, both terms include the factor *a cos*(2*πkn*/*N*). Since the cosine function is even (cos(−θ) = cos(θ)), the cosine terms for *k* and *N* − *k* are identical and add together, reinforcing the real component of the IFT. Conversely, the sine terms possess differing signs. The term from *X*[*k*] includes:

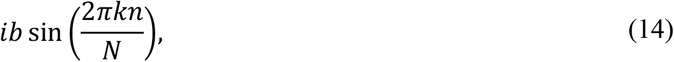

while the contribution from *X*[*N* − *k*] comprises:

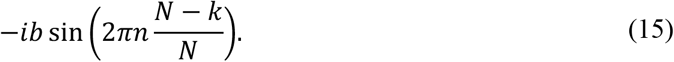

Due to sine function odd property (sin(−θ) = − sin(θ)) these sine terms negate each other. Therefore, both terms include only the real factor −*b* sin(2*πkn*/*N*). Hence, adding up the remaining terms from conjugate pairs results in a real value in the form of:

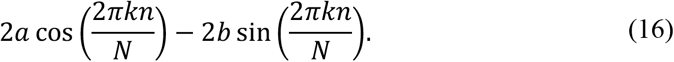

This proof solidifies the importance of maintaining conjugate symmetry when manipulating CCs in Fourier analysis of gene expression data, ensuring that the numerical transformations preserve the original data real integrity and characteristics.

### Conjugate pair identification for even and odd length gene expression profiles

A fundamental aspect of applying modifications to CCs involves accurately identifying the indices of conjugate pairs, which varies depending on whether the profile length (number of detected genes, N) is even or odd. For even-length profiles (where *N* is even), the Nyquist component (*X*[*N*/2]; indicating the upper limit of complex content) lacks a distinct conjugate pair, i.e. a self-conjugate component. For the remaining components, specifically for *X*[*k*] where 1 ≤ *k* < *N*/2, the conjugate pair is identified as *X*[*N* − *k*]. It is imperative to modify *X*[*k*] and corresponding adjust *X*[*N* − *k*] to uphold conjugate symmetry. For example, in a profile with *N* = 6, *X*[1] and *X*[5] form conjugate pairs, as do *X*[2] and *X*[4], while *X*[3] is self-conjugate.

In the case of odd-length profiles (where *N* is odd), for any component *X*[*k*] where 1 ≤ *k* ≤ (*N* − 1)/2, the conjugate pair is *X*[*N* − *k*]. Modifications to *X*[*k*] must be mirrored in *X*[*N* − *k*] to ensure the preservation of conjugate symmetry. For instance, with *N* = 5, *X*[1] and *X*[4] are conjugate pairs, just as *X*[2] and *X*[3]. For an odd-length profile, there is no exact Nyquist component.

Regardless of the profile length, the 0-th component, Direct Current (DC) component, represents the overall average or baseline level of the signal:

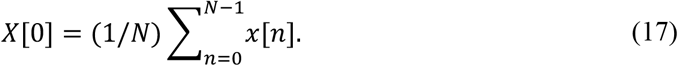

Given its representation of the baseline transcriptional activity, modifications to this component can disproportionately skew the entire gene expression profile upon transformation back to the original data space. To preserve the intrinsic profile baseline and ensure that synthetic profiles maintain a realistic representation of overall gene expression levels, the DC component is kept unaltered.

### Implementation steps for generating synthetic single-cell

Synthetic cell generation commences with the transformation of cell gene expression profiles into complex space using DFT. Next, CCs are stochastically selected for modification. Modification factors are sampled from pre-defined amplitude distributions associated with each component, aggregated across cell groups with similar expression profile (such as cell-type or cluster of cells). These factors are applied symmetrically to both the sampled CCs and their corresponding conjugate pairs to ensure the real-valued nature of the profile is preserved. Subsequently, the IFT is employed to construct synthetic gene expression profiles. This process is mathematically formalized as follows:

Given a population of cells, for each cell, the DFT is applied to its gene expression profile, *x*_*j*_[n], yielding complex components *X*_*j*_[*k*], where *n* indexes genes, *k* indexes complex component, and *j* indexes cells:

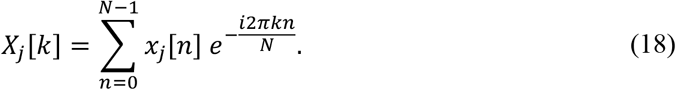

Next, for each complex component *k*, the mean *μ*_%_ and variance 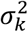 of its amplitude are calculated across the cell group:

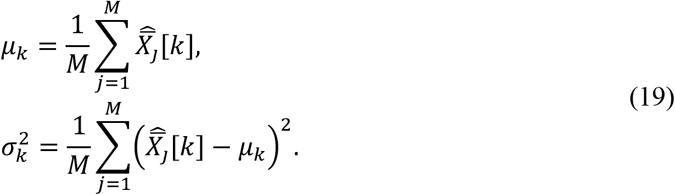

Here, *M* denotes the total number of cells in the group and 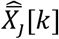 denotes the amplitude of the *k*-th complex component for the *j*-th cell, 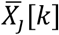, scaled by its Euclidean norm (*L*2 norm) in the form of:

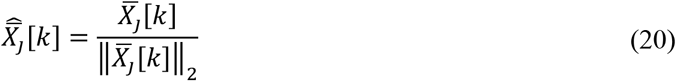

and

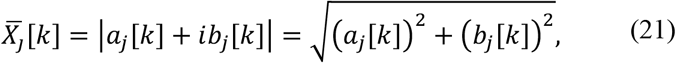

respectively.

A Gaussian distribution is then defined for each complex component *k*, informed by the aggregate statistics, from which the modification factor *A*_%_ is sampled:

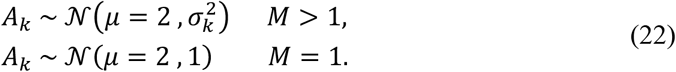

The Gaussian distribution is utilized for its symmetric and unbiased nature, ensuring an even distribution of both enhancement and attenuation modification coefficients. The choice of centering the distribution at 2 directly corresponds to the propagation modification term, (*A* − 1)*X*[*m*]*e*^*i*2*πmn*/*N*^, as shown in equation (6). If the group contains only one cell (*M* = 1), the modification factor is sampled from a Normal Gaussian distribution with its mean shifted by +2 units. Therefore, this method accounts for modifications applied to both individual cells and groups of cells during the synthesis process.

We note three key aspects: (1) Each CC encapsulates a unique variation pattern that, despite varying in magnitude and phase across cells, captures modes of variation present in groups of cells with similar gene expression profiles, such as specific cell types or clusters. Equation (22) has been formulated to mathematically embody this concept. (2) To ensure proportional expansion across different cell populations, all pre-defined cell groups are uniformly scaled according to the desired expansion magnitude. This approach maintains the relative proportions among cell groups, ensuring that the expanded populations accurately reflect the original abundance distribution. (3) For a fully stochastic synthesis process, cells are randomly selected for synthesis within each group.

The sampled modification factor is then applied to randomly selected *k*-th complex component and its conjugate pair:

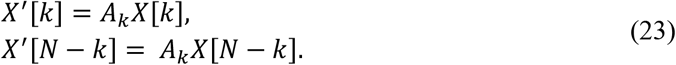

Finally, the adjusted complex components, *X*’, are transformed back into the gene expression space to construct the synthetic profile for a new cell:

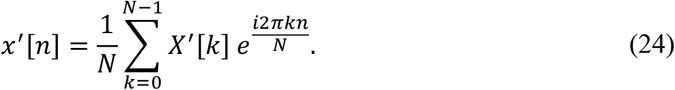

Overall, this representation formalizes the generation of synthetic scRNA-seq data by introducing controlled and fine-grained variations, thereby simulating the natural biological variability inherent in authentic datasets. At the core of this method is a mathematical foundation that modulates the degree of deviation from the original gene expression profiles by altering the number of CCs subject to modification. This capability provides a systematic method for adjusting the fluctuation introduced into synthetic profiles.

### Pre-synthesis processing

Two preliminary processing steps are undertaken prior to synthesis. (1) Unique molecular identifier (UMI) counts for each cell are scaled to counts per million and then transformed using the natural logarithm to stabilize expression variability across cells. (2) The synthesis procedure is selectively applied only to the most variably expressed genes. These genes, manifesting biological variation indicative of distinct cellular states or phenotypes, are targeted to mitigate the amplification of stochastic noise prevalent in genes with lower variability.

### Post-synthesis processing

Three processing steps are undertaken after synthesis. (1) To enhance the biological plausibility of the synthesized gene expression profiles and mitigate potential noise introduced during the IFT, a smoothing step is implemented. This step leverages the original gene expression profiles and their nearest neighbors, as defined by the nearest neighbor (NN) graph. For each synthesized cell, gene expression values are averaged with those of its original counterpart and nearest neighbors. (2) Since UMIs represent counts of sequences, negative values are biologically implausible. To address any negative values that might arise during the transformation process, a Rectified Linear Unit (ReLU) activation function is applied to each component of the synthesized profile, ensuring all values remain non-negative:

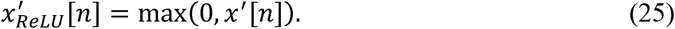

(3) The synthesized raw counts are derived from their original raw counterparts by taking into account the relative change rate between the synthesized data and their original normalized counterparts. This completes the synthesis process as delineated by the scGFT Fourier transform-based generative model (**Fig. S2**).

### Assessing the deviation of synthesized gene expression profiles from original cells

To evaluate the fidelity of synthetic gene expression profiles in mimicking the transcriptional characteristics of original cells, the deviations between the expression profiles of original cells and their synthesized counterparts is quantified.

Let *O* = (*o*_1_, *o*_2_, …, *o*_*N*_) represent the expression profile of an original cell, where *N* is the total number of genes and *o*_*g*_ indicates the expression level of gene *g* in the original cell. Similarly, let 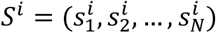 denote the gene expression profile of the *i*-th synthesized cell derived from *O*, with *i* ϵ {1,2, …, *L*} representing the index of synthesized cells, where *L* is the total number of synthesized cells derived from *O*. Here, 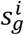 indicates the expression level of gene *g* in the *i*-th synthesized cell. Then, the relative difference in expression for gene *g* between the original cell and each synthesized cell is computed as:

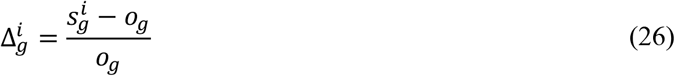

The average deviation per gene across all synthesized cells is then calculated as:

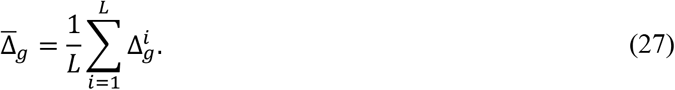

This measure provides an indication of how closely the synthetic cells gene expression profiles resemble that of the original cell, thus assessing the impact of synthesize process on the original cell transcriptional landscape.

### Single cell RNA-seq data processing

For the processing of PRJEB44878 scRNA-seq data, we utilized the CellBridge workflow^28^, starting from FASTQ files. Canonical QC parameters were employed throughout the processing steps, including the minimum number of UMIs per cell (set to 750), the minimum number of genes per cell (set to 250), the minimum number of cells expressing a given gene (set to 3), the percentage of mitochondrial genes per cell (set to 15), and the threshold used for doublet identification (set to 0.25). Default parameters were used unless otherwise specified.

We adhered to the standard Seurat^23^ (v5.0.1) pipeline for scRNA-seq data processing. Data normalization was first carried out sing the “NormalizeData” function with the “LogNormalize” method and a scale factor of 10^>^. This was followed by identifying the top 2,000 variable genes using “FindVariableFeatures”. The data were then centered and scaled using “ScaleData”. Principal Component Analysis (PCA) was executed with “RunPCA”. To adjust for batch effects in the experimental data, the Harmony^29^ (v1.2.0) algorithm was applied via “RunHarmony” on PCA-reduced data, considering sample-specific variations. Neighbors were identified using “FindNeighbors” based on the harmonized components, which facilitated the clustering of cells into distinct groups with “FindClusters” using a resolution of 0.7. The combined dataset of original and synthetic cells underwent another round of features identification, scaling, PCA, batch correction (considering sample-specific variations), neighbor identification, and clustering. Finally, we visualized the integrated data landscape using “RunUMAP”. Default parameters were used, unless otherwise specified.

In the simulated data processing, cells with fewer than 250 detected genes and genes expressed in fewer than 500 cells were excluded. Normalization (counts per million) was performed, and highly variable genes (n=2,000) were identified, followed by scaling, PCA, and neighbor identification, using the standard Seurat pipeline. The combined dataset of original and synthetic cells underwent another round of features identification, scaling, and PCA. For clustering the integrated data, the k-means approach was adopted to align the number of clusters with the original dataset, ensuring a conservative comparison by equating the grounds. This process was executed using the “kmeans” function from the “stats” package (v3.6.2), applied to the PCA embeddings. Default parameters were used, unless otherwise specified.

### Construction of cell-type specific networks and module identification

In constructing cell-type specific networks, methodical steps were undertaken to refine and prepare the data according to Genix protocols^30^. Genes expressed in less than 25% of cells and cells expressing fewer than 1% of genes were removed. Subsequently, the count expression matrix underwent normalization to the mean UMI library size, scaling each cell’s expression sums to the mean total counts. Finally, networks were inferred, and modules were identified using the “constructNets” and “compileNets” functions, respectively, from the Genix R package (v1.0.0). Default parameters were used, unless otherwise specified. In this scenario, networks were inferred for AT0, AT1, and AT2 epithelial subtypes.

We devised a method to merge AT subtype-specific modules, thus distinguishing between general modules (shared across all ATs) and more specific ones. Employing the Jaccard Index, we measured the similarity among the modules, generating a square matrix where rows and columns correspond to each module. We then applied hierarchical clustering to identify clusters of modules with similar gene compositions, thereby redefining “module” to refer to these merged (unified) clusters. To determine the optimal number of clusters, we utilized silhouette analysis (“clsuter” R package, v2.1.4), which measures how similar each point in one cluster is to points in neighboring clusters. By computing the average silhouette width for a range of cluster numbers up to 50, we identified the number of clusters that maximizes this metric, indicating the best separation between clusters. Default parameters were used, unless otherwise specified.

### Quantification of gene module associations with biological states

To identify modules that are most predictive of biological states (a feature selection task), we employed a regularized regression approach using LASSO (least absolute shrinkage and selection operator). We utilized the binomial configuration of the regression model in the “cv.glmnet” function from the glmnet R package (v4.1.8) to quantify module associations with the biological state under study versus a baseline state (e.g., diseased versus healthy or cell type X versus cell type Y). During the fitting procedure, the penalized regression model optimizes the coefficients to minimize a penalized loss function that combines the goodness of fit with a penalty for large coefficients. The resulting optimized coefficients indicate the significance of each module in differentiating between two conditions. Specifically, a positive coefficient implies that higher expression of a module is associated with the state under study, while a negative coefficient implies an association with the baseline state. In this scenario, cell type AT0 is considered the baseline, while AT1 and AT2 are the states under study.

To assess the statistical significance of the observed coefficients, we implemented a bootstrap approach. For each biological state, we performed 5,000 bootstrap replicates of the model and retrieved the coefficients for each module. We then generated null distributions for each module under the null hypothesis that there is no association (i.e., the true coefficients are zero). This involved simulating normally distributed values with a Gaussian distribution mean of zero and a standard deviation matching that of the observed bootstrap coefficients. P-values for the observed coefficients were computed by comparing them against their respective null distributions using a two-tailed empirical approach. This was achieved by calculating the minimum proportion of null distribution values greater or smaller than the observed coefficient, multiplied by two to account for both tails.

### Additional data analysis and visualization functions

Differentially expressed genes were identified using the “FindMarkers” function from the Seurat R package (version 4.3.0) with the Wilcoxon rank sum test. Genes with an absolute log2-fold change larger than 1 and an adjusted p-value less than 0.05 were considered significant. The “DotPlot” function from Seurat was used to visualize gene expression with dot plots. The “DimPlot” function was used to visualize the final UMAP on a 2D scatter plot. The “AddModuleScore” function calculated module scores for each cell based on the average expression of associated gene lists. Enrichment between inferred modules and reference Gene Ontology (GO) pathways was analyzed using the “newGeneOverlap” function from the GeneOverlap R package (version 1.26.0). Plots were created using the ggplot2 R package (version 3.3.5) unless otherwise noted. Default parameters were used, unless otherwise specified. All analyses were performed using R version 4.3.2.

## Data availability

The lung organoid FASTQ files are available in the European Nucleotide Archive database under accession code PRJEB44878. The PRJEB44878 scRNA-seq count matrix and metadata have been deposited at Zenodo repository and are publicly available under record no. 11166226. The lung distal airways scRNA-seq data is publicly available at NCBI Gene Expression Omnibus, accession numbers GSE178360. The simulation data have been deposited at Zenodo repository and are publicly available under record nos. 11166226 and 11165521.

## Code availability

The scGFT source code (R package) is available on GitHub at https://github.com/Sanofi-Public/PMCB-scGFT.

## Acknowledgments

We want to express our gratitude to Dr. B. Plaster, Professor of Physics at the Department of Physics and Astronomy, University of Kentucky, Dr. S. Kleinstein, Professor of Pathology, Dept. of Immunobiology, Yale School of Medicine, as well as Dr. H. Mattoo, Dr. G. Gaglia and Dr. M. T. Valerius, Sanofi Precision Medicine and Computational Biology (PMCB) for their valuable comments on the manuscript. We express our gratitude to Dr. V. Savova and Dr. E. de Rinaldis, Sanofi PMCB, for their support and strategic direction. We are grateful for the use of the Sanofi R&D compute platform, Magellan, for facilitating data analysis. This work was supported by Sanofi US.

## Supplementary Figures

**Figure S1.**
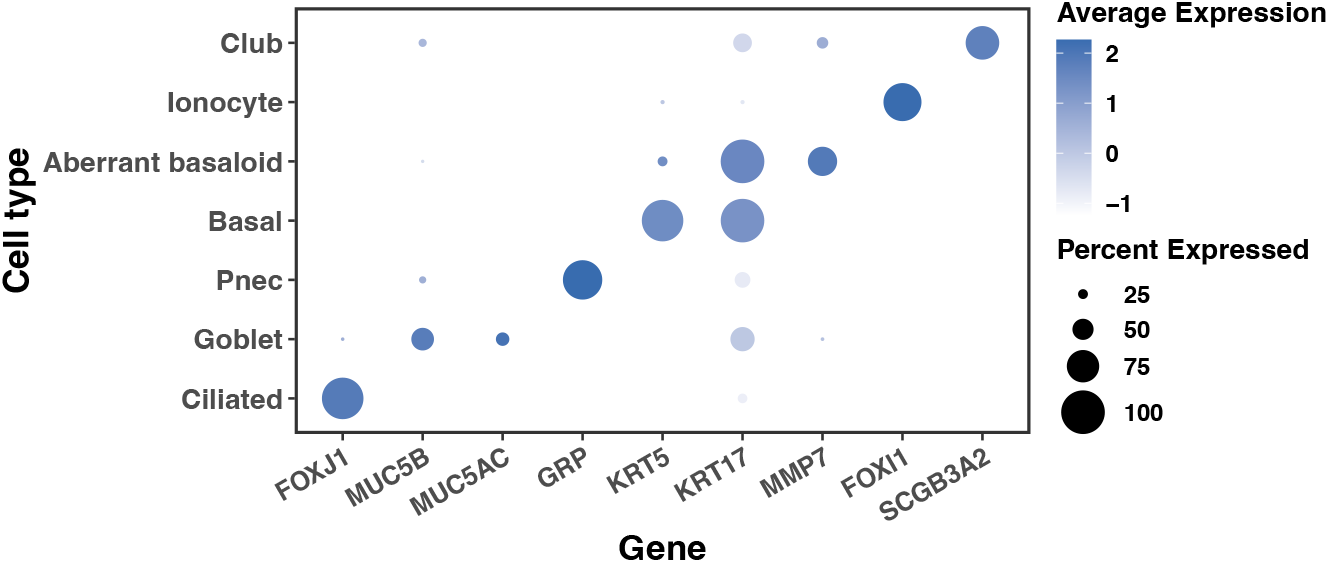
Expression levels of epithelial subtype markers. Dot-plot displaying selected marker genes for each of the different subtypes of epithelial cells in the PRJEB44878 dataset. The color of each circle represents the average expression level of the gene, while its size indicates the percentage of cells expressing that particular gene.

**Figure S2.**
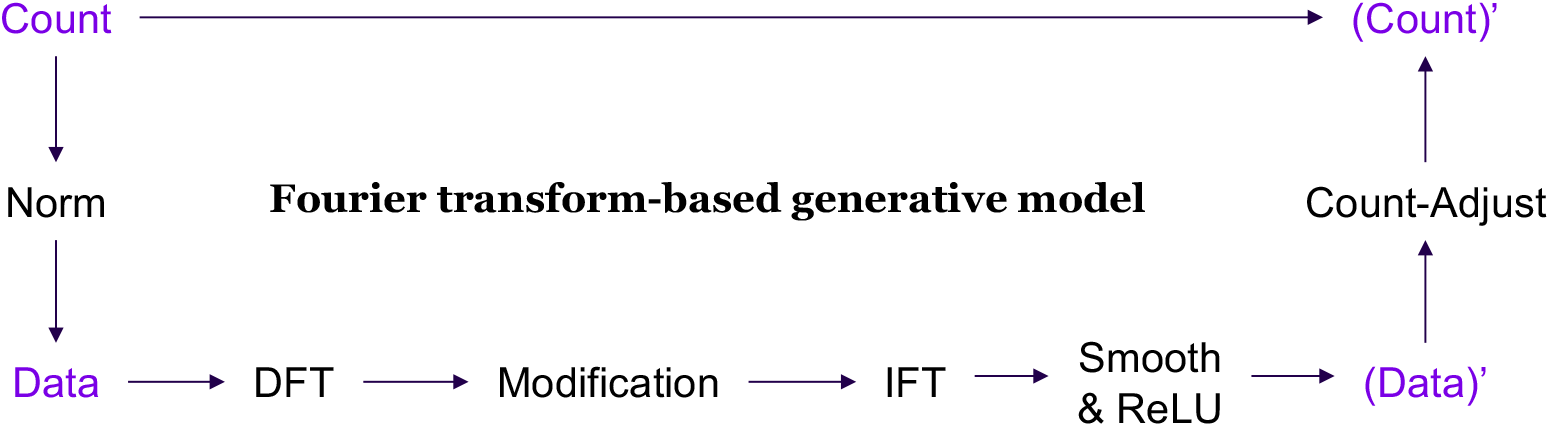
Overview of the scGFT Fourier transform-based generative model. The process commences with the normalization of raw counts, followed by the application of the Discrete Fourier Transform (DFT) to extract complex components. These components are then modified and subjected to an Inverse Fourier Transform (IFT) to map back to the original data space. To mitigate potential noise introduced during the IFT, a smoothing procedure is implemented, followed by a Rectified Linear Unit (ReLU) activation function to clip any residual negative values generated during the IFT process to zero. Finally, the synthesized raw counts are derived from their original raw counterparts by taking into account the relative change rate between the synthesized data and their original normalized counterparts.

